# Empirical patterns of environmental variation favor the evolution of adaptive transgenerational plasticity

**DOI:** 10.1101/426007

**Authors:** Jack M. Colicchio, Jacob Herman

**Affiliations:** Department of Plant and Microbial Biology, University of California Berkeley, Berkeley, CA, USA; Department of Organismic and Evolutionary Biology, Harvard University, Cambridge, MA, USA

**Author notes:** Corresponding Author: Jack Colicchio, Koshland Hall 361, Berkeley, CA, 94720.

**Keywords:** Transgenerational plasticity, epigenetics, local adaptation, climatic oscillations

## Abstract

Effects of parental environment on offspring traits have been well known for decades. Interest in this transgenerational form of phenotypic plasticity has recently surged due to advances in our understanding of its mechanistic basis. Theoretical research has simultaneously advanced by predicting the environmental conditions that should favor the adaptive evolution of transgenerational plasticity. Yet whether such conditions actually exist in nature remains largely unexplored. Here, using long-term climate data, we modeled optimal levels of transgenerational plasticity for an organism with a one-year life cycle at a spatial resolution of 4km2 across the continental US. Both annual temperature and precipitation levels were often autocorrelated, but the strength and direction of these autocorrelations varied considerably across the continental US and even among nearby sites. When present, such environmental autocorrelations render offspring environments statistically predictable based on the parental environment, a key condition for the adaptive evolution of transgenerational plasticity. Results of our optimality models were consistent with this prediction: high levels of transgenerational plasticity were favored at sites with strong environmental autocorrelations, and little-to-no transgenerational plasticity was favored at sites with weak or non-existent autocorrelations. These results are among the first to show that natural patterns of environmental variation favor the evolution of adaptive transgenerational plasticity. Furthermore, these findings suggest that transgenerational plasticity is highly variable in nature, depending on site-specific patterns of environmental variation.

## Introduction

Natural selection can produce adaptation only if the selective environment is reliably encountered over generations, or in other words, if selective environments are statistically predictable. Early models of evolution envisioned fitness landscapes that were static, such that populations adapt over the course of generations to one or another environment (Fisher 1930). While this form of adaptation optimizes phenotypes in homogenous environments, the more realistic scenario of environmental heterogeneity in both space and time limits the adaptive value of such constitutive genetic expression (Sultan 2015. In variable environments, the capacity to modify phenotypes in response to predictive environmental cues allows organisms to match their traits to the specific patch of habitat in which they find themselves, a phenomenon termed adaptive within-generation plasticity (Ghalambor et al. 2007; Nicotra et al. 2010). Investigating the predictability of environmental cues in nature is therefore a major research goal in ecology and evolution.

Over the last three decades, it has become clear that effects of parental environments on offspring phenotypes (i.e., *transgenerational plasticity*) are remarkably common (reviewed by (Mousseau and Fox 1998; Uller 2008; Bonduriansky and Day 2009; Holeski et al. 2012; Conrath et al. 2015; Sultan 2015). For instance, when *Mimulus guttatus* plants experience herbivory, their offspring increase production of defensive leaf trichomes (Holeski 2007; Colicchio et al. 2015; Colicchio 2017). Similarly, when the aquatic crustacean *Daphnia cuculatta* senses predator cues, it produces offspring with a defensive ‘helmet’ that protects against predation by midge larvae and cladocerans (Agrawal et al. 1999). Such inherited environmental effects can be transmitted from parent to offspring (and to additional generations in some cases) by diverse mechanisms, including heritable epigenetic modifications (i.e., DNA methylation marks, histone modifications, and small RNAs) and the allocation of nutritive resources, hormones, mRNAs, and regulatory proteins to seeds or eggs (these mechanisms are not mutually exclusive; (Herman and Sultan 2011; Jablonka 2013). As more research has focused on transgenerational plasticity, it has become clear that these effects are highly variable (Herman and Sultan 2016; Colicchio 2017; Groot et al. 2017) and nearly absent in some cases (Ganguly et al. 2017). Empirical investigations in diverse plant and animal systems have confirmed that transgenerational environmental effects can be adaptive when parent and progeny environments match (i.e., under positive intergenerational environmental autocorrelations; see e.g., Bilichak et al. 2012; Herman et al. 2012; Rasmann et al. 2012; Slaughter et al. 2012; Verhoeven and van Gurp 2012; Dantzer et al. 2013; Lopez Sanchez et al. 2016; Walsh et al. 2016; Wibowo et al. 2016). Further, Dey et al. (2016), Graham et al (2014)., and Sikkink et al. (2014) have demonstrated transgenerational plasticity can evolve in experimental settings.

These results motivated evolutionary research probing the theoretical scenarios in which transgenerational plasticity is expected to evolve adaptively. A central insight is that natural selection should favor specific forms of plasticity depending on the precise patterns of environmental variation experienced by a population (Shea et al. 2011; Sultan and Spencer 2002). Existing theory on the evolution of both within-generation (Tufto 2015; reviewed by Scheiner 1993; Schlichting and Pigliucci 1998) and transgenerational plasticity (Lachmann and Jablonka 1996; Räsänen and Kruuk 2007; Kuijper et al. 2014; Prizak et al. 2014; Leimar and McNamara 2015) has demonstrated that plasticity can evolve when environmental conditions are correlated across time, there is little to no cost of responding to environmental cues, and there is genetic variation in reaction norm slope.

Transgenerational plasticity in particular is likely to evolve when parental and offspring conditions are either positively or negatively correlated (Proulx and Teotonio 2017), with the magnitude of the correlation being the primary factor determining the optimal level of transgenerational plasticity. Recent models have shed light into the evolution of transgenerational plasticity in patchy environments (Leimar and McNamara 2015), explicitly testing the conditions that favor deterministic vs. randomizing maternal effects (Proulx and Teotonio 2017), how migration and population structure impact the evolution of transgenerational plasticity (Greenspoon and Spencer 2018), the optimal levels of epigenetic resetting between generations (Uller, English, and Pen 2015), and the interaction between the evolution of within-generation and transgenerational phenotypic plasticity (Kuijper and Hoyle 2015). Additionally, other groups have developed systems comparing invasion probabilities of lines with various epigenetic modifier loci (Furrow and Feldman 2014) and applied information theory (Donaldson-Matasci, Bergstrom, and Lachmann 2013) to the evolution of transgenerational phenotypic plasticity.

As formally shown through a variety of models, when environmental autocorrelations increase, the optimal degree of transgenerational response also increases (e.g., McNamara et al. 2016). In other words, for transgenerational plasticity to be adaptive, the environment must not only be variable but also predictable (Burgess and Marshall, Oikos, 2014) from one generation to the next. The scale of environmental variation can also be described in terms of environmental grain (Gillespie 1974), where the relative “coarseness” describes whether the environment fluctuates rapidly or slowly between states. When the environmental grain is too coarse, genetic adaptation is expected to predominate over forms of plasticity (Banta et al. 2007). When the coarseness is too fine-grained, transgenerational plasticity is not expected to evolve because the environmental information sensed by the parent is out of date when progeny receive it (McNamara et al. 2016). In the case of organisms with relatively fixed generation times, the autocorrelation between parental environmental cues and offspring selective environments provides a simple quantification of the levels of transgenerational plasticity that should maximize the mutual information between phenotype and environment. A common theme across all of the theoretical literature is that these autocorrelations are likely the most important factor in the adaptive evolution of transgenerational plasticity (Burgess and Marshall, 2014). This consensus motivated us to assess the presence of autocorrelations across this scale of environmental grain.

Despite this surge of experimental evidence, molecular understanding, and theoretical interest, no study to date has examined long-term environmental data for the presence of such environmental autocorrelations. Although evolutionary research has traditionally focused on how the average environmental conditions differ across a landscape, there is no reason to expect that the scale and predictability of environmental variation is any less complex or ubiquitous than variation in mean environmental conditions. As prior modelling studies have demonstrated (Uller et al. 2015), the *spatial variation* in the *temporal predictability* of environmental variation is expected to drive the evolution of transgenerational effects across a heterogeneous landscape.

In this study, we test if empirical patterns of climatic variation allow for the evolution of within-generation plasticity, transgenerational plasticity, and multigenerational epigenetic inheritance across different local climate regimes. We use 120 years of fine-scale (4km2) climate data spanning the coterminous U.S. to test for auto- and cross-correlations in temperature and precipitation levels across years. We found many significant correlations that vary widely in both magnitude and direction across the US. We then constructed separate models, with summer annual plants in mind, for temperature and precipitation to determine the degree of transgenerational plasticity that would maximize fitness in each of these sites across the U.S. Furthermore, by running each model using raw environmental data and the residuals after removing the effects of directional climate change, we were able to inspect how climate change alters the benefits associated with transgenerational plasticity. These results allow us to detect where transgenerational plasticity is expected to evolve given patterns of environmental variation over the past 120 years.

In our precipitation model, we examine transgenerational effects that persist for up to three generations (Figure 1a), as multiple experimental studies have found that environmentally induced epigenetic and phenotypic effects can persist for at least this long (e.g., Whittle et al. 2009; Akkerman et al. 2016), and in some cases for far longer (Vastenhouw et al. 2006; Rechavi et al. 2011). While we do not consider specific mechanisms of transgenerational plasticity, prior work has demonstrated that the offspring of plants exposed to drought stress have higher survival in drought conditions, partially mediated through enhanced root growth phenotypically and altered DNA methylation patterns at the molecular level (Herman and Sultan 2016). In our temperature model (Figure 1b), we also determine the degree of within-generation plasticity that would maximize fitness, in response to both early and late-season temperatures. In plants, transgenerational effects of temperature have primarily been demonstrated to shift phenology such as flowering time (Case et al. 1996) and dormancy (Chen et al. 2014), but other phenotypes such as rosette diameter in Arabidopsis are also impacted by parent temperature (Groot et al. 2017). In animals, egg size, survival, developmental rate, melanisation, and heat-shock survival were all shown to be impacted by parent temperature (*see review*: Donnelson et al. 2017). DNA methylation likely contributes to transgenerational effects of temperature, and small RNAs also appear to be a major contributor (Houri-Zeevi and Rechavi 2017).

**Figure 1:**
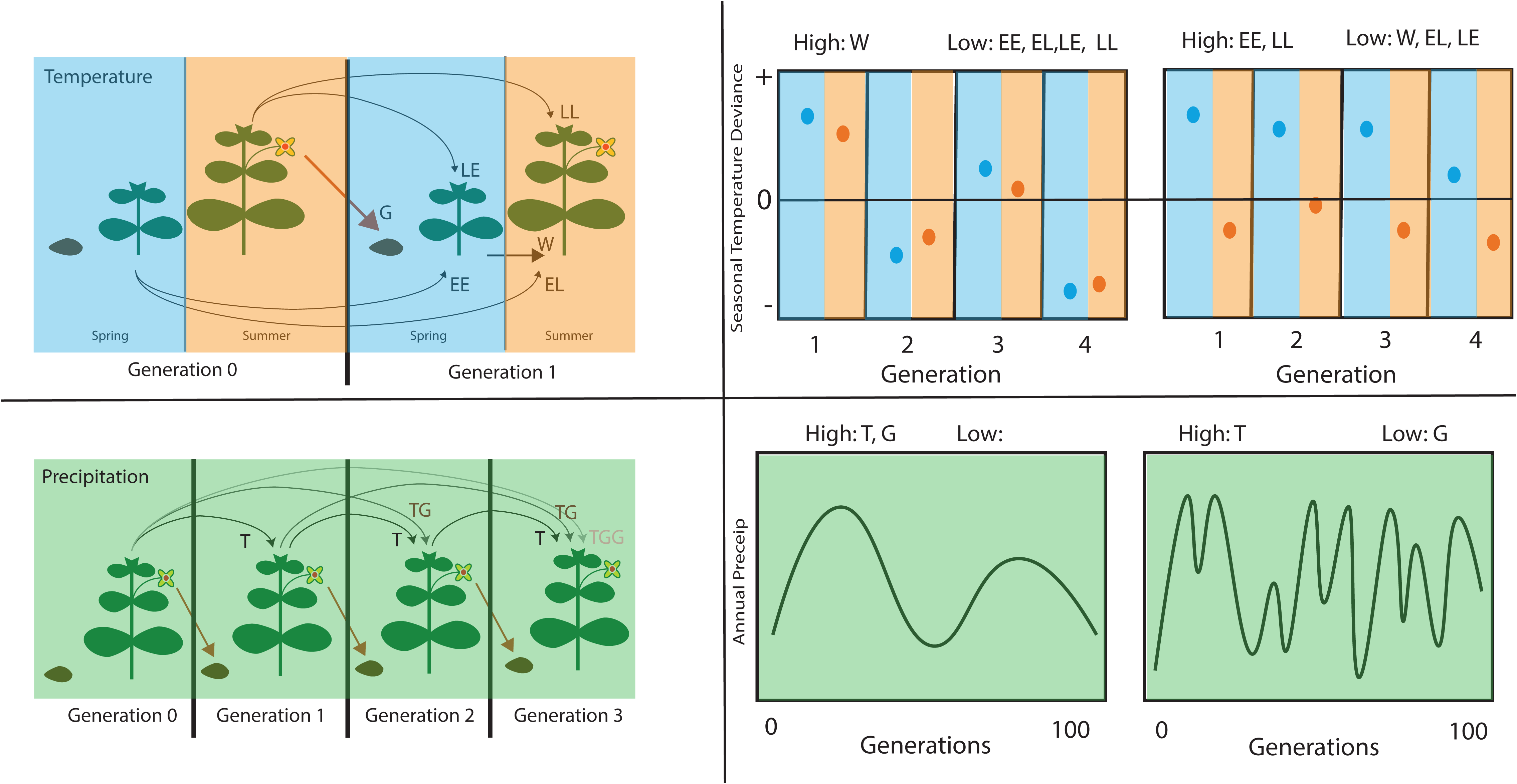
Schematic depicting the ecological motivations (summer annual plants) and theoretical underpinnings for the evolutionary modeling of plasticity traits (A and C), and the types of environmental fluctuations that may influence their evolution. (A) Temperature plasticity model. In the abbreviations, **E** denotes the Early growing season (spring) and **L** denotes the late growing season (summer). The first letter represents the relevant season during the parental generation and the second letter represents the relevant season in the offspring generation (e.g., EL denotes effects of parental early growing season temperature on offspring phenotypes late in the growing season). Within generation developmental changes in response to early season environment (W) are also considered in this model. Additionally, the long-term average environmental conditions at a specific area determine the genetic baseline phenotype of an individual (G). (B) On the left we see an example of an environment with high within season autocorrelations for temperature (hot springs tend to be followed by hot summers), but low inter-annual autocorrelations (a hot year does not tend to be followed by another hot year) that selects for within generation plasticity but not transgenerational plasticity. On the right, a situation where spring and summer temperatures are not correlated with each other, but we do find that environmental oscillations lead to a string of warmer than average springs and cooler than average summers, in this situation transgenerational plasticity (EE and LL) but not within generation plasticity is expected to be optimal. (C) Precipitation plasticity model. The amount of precipitation experienced by an individual can lead to transgenerational effects in the next generation (T), as well as persist for two (TG) or three (TGG) generations. (D) On the left, relatively gradual decadal oscillations give value to transgenerational effects that persist for multiple seasons (T, TG, and TGG). On the right shorter period climatic oscillations may favor parental effects (T), but not multi-generation effects (TG or TGG).

Our optimality models extend previous models (e.g., Kuijper and Hoyle 2015; Leimar and McNamara 2015), by considering multiple different seasonal timepoints in the parental and offspring generations in which deterministic transgenerational effects can be induced and alter phenotypes. We also quantitatively simplify prior methods to allow these multiple parameters to be considered across hundreds of thousands of locations. We compare geometric mean fitness across the 120 years of climatic data for individuals that utilize different classes of parental information to different degrees. This approach is similar to how Proulx and Teotonio (2017) used geometric mean fitness to compare invasion success in individuals exhibiting a variety of different maternal effect strategies. Rather than assigning a formal genomic framework to our data, we consider a theoretical scenario in which there is no sexual reproduction, or gene transfer of any kind, and where alleles altering transgenerational plasticity can vary in magnitude and direction. By distilling down our models to identify the optimal values of transgenerational plasticity at a given site, we recapitulate the finding from more dynamic models that plasticity is tied directly to environmental autocorrelations and are able to apply these theoretical findings to real world climate data. These results suggest that climatic factors could be sufficient to select for locally adaptive variation in transgenerational plasticity across the landscape. Finally, biologists can use these findings to design experiments by identifying areas of the U.S. where transgenerational effects are more apt to evolve.

## Methods

### Descriptive statistics

Mean monthly temperature and precipitation at a 4km resolution from 1895-2014 (LT81m) were downloaded from the PRISM climate group web server (PRISM Climate Group, 2004). In short, PRISM uses climate averages from between 1981-2010 as a predictor grid, and then utilizes station networks with at least 20 years of data to model monthly temperature and precipitation across the US. The emphasis on this dataset is long-term consistency making it ideal for our purposes. Individual yearly values were concatenated using the QGIS merge raster function (Quantum GIS geographic information system 2012) to create a single data frame, and exported in the .RData format for downstream analysis. For precipitation data, October was chosen to represent the start of the “hydrologic” year in order to more accurately capture water availability patterns during the growing season. For temperature data, mean daily maximum temperature was calculated for March-May as a measure of early growing season temperature for a given year, and July-September mean daily maximum temperature for late growing season temperature. Autocorrelations were calculated at lags between 1 and 12 years (i.e., environmental correlations were calculated between year X and year X+1, year X and year X+2…year X and year X+12).

### General modeling framework

Mathematical models were constructed in R for both precipitation and temperature patterns to compare how individuals that use within-generation plasticity, transgenerational plasticity, and genetic inheritance to varying degrees differ in their capacity to match their phenotype with the environmentally optimal phenotype for a given year. In these models, there are hundreds (precipitation models) or thousands (temperature models) of competing genotypes, each representing unique points of parameter space for alleles that modify the extent to which environment affects phenotype. Trait value is a measure of the expected environment (temperature or precipitation) and is determined by a combination of the mean environment at a given site over all years, and terms that modify this value based on recent environmental information. Each genotype is in essence a climatologist, that utilizes genetic information (based on mean precipitation over the 120 years at a site), transgenerational plasticity, or within generation plasticity (only in temperature model) to come up with an expected environment that it will face. This expected environmental value is equivalent to a phenotype, and the closer this phenotype is to the actual environment experienced, the higher the fitness that genotype will have for a given generation.

While this framework is identical for precipitation and temperature modeling, inherent differences in precipitation and temperature variables lead to us considering a different set of parameters for each variable, allowing us to ask related but unique questions regarding transgenerational inheritance. Precipitation can accumulate as snowpack, bodies of water, or soil moisture, such that the cumulative precipitation over the course of the water year will determine to a large extent the amount of water available to a plant. On the other hand, the effects of temperature are much more immediate and transient, such that a particularly cold spring will not “keep the plant cool” over the summer, in the way that a particularly wet spring could provide moisture during a summer of drought. For this reason, we decided to extend our temperature models to compare patterns across different segments of the growing season, and different forms of plasticity both within and between a single generation. For precipitation we only considered annual hydrologic year precipitation without breaking it down by seasons, but we did consider the possibility of multigeneration persistence of transgenerational plasticity.

For each locale, mean annual precipitation (or temperature) across the 120 years (***<u>P</u>***) is calculated, and this statistic is used as the baseline phenotype of all genotypes in the raw data variant of the model (Appendix 1). In the residual variant of this formula, a linear regression was fit over the time-course, and residuals were used as the climate values for each year, with a baseline phenotype of 0. This baseline phenotype is then be adjusted to varying degrees by the parent environment, such that different genotypes will weigh the contribution of parent environment to a different extent. For both the precipitation and temperature model we calculated the phenotypes produced by each of 176 (precipitation) and 3,125 (temperature) genotypes at each site (481,631), for each year (119). Then by comparing the phenotype produced during a season with the actual environment of that season, we imposed a linear cost on fitness based on the distance between phenotype and the actual temperature or precipitation of that year (Appendix 1). We then the calculated geometric mean fitness of every genotype at each site independently to predict which genotype would have the greatest increase in frequency over the course of the time series, and we considered this the optimal phenotype for that site.

This modeling framework represents a variant of other transgenerational plasticity models where the direct parent environment alters offspring phenotype. We model the transgenerational effect as a linear reaction norm with slope *m* with respect to the environment experienced at a particular previous point in time (for the precipitation model this represents the water year experienced by the past generation, or in the temperature model the temperature experienced in the current generation’s spring, the previous generation’s spring, or the previous generation’s fall). In the case of multigenerational effects, our *g* terms linearly reduce the norm of reaction slope of the grandparental and great-grandparental generation relative to the effects of the parental generation (Appendix 1). This approach is similar to the analytical models designed by Uller, English, and Pen (2015) where maternal effects were modeled as a “linear reaction norm with respect to the mother’s *perceived* environment” where the perceived environment was the environmental state of the previous generation with an additional normally distributed error term. Leimar and McNamara (2015) utilize a more complex model where adult phenotype is modeled as a logistic function wherein the *a*_*ma*t_ term determines the weighting of maternal environmental cue, as well as two terms (*m*_*mat*_ and *d*_*mat*_) that control the weighting of maternal phenotype transgenerational plasticity and direct parent environment transgenerational plasticity. Proulx and Teotonio (2017) consider six different classes of inheritance strategies competing in environments that switch between two states with variable frequencies. In their modeling framework the strategies aDME and mDME correspond to two-state variants of positive (m>0) and negative (m<0) transgenerational effects, respectively, as modeled here. Finally, Kuijper and Hoyle (2015) model maternal effects as a linear transgenerational reaction norm but on parental phenotype rather than parental environment. In our models, we consider fitness to decrease linearly as an individual’s phenotype moves further from the phenotypic optimum at a point in time. We compare the geometric mean fitness of individuals expressing different strategies over the 120 years to find the strategy most likely to invade. Similar to most previous models (Uller, English, and Pen 2015; Proulx and Teotonio 2017; Lachmann and Jablonka 1996; but see Leimar and McNamara 2015; Greenspoon and Spencer 2018), we base our model on haploid asexually reproducing individuals. Our temperature model extends previous work by explicitly breaking down both the life cycle of the parent and offspring generations between early and late growing season, allowing for five different temporal classes of plasticity (four types of transgenerational plasticity, and within-generation plasticity).

## Results

Both mean annual precipitation and growing season temperatures vary immensely across the US (Figure 2 and S1), but for the evolution of locally adaptive phenotypic plasticity, it is the patterns of variation that are more relevant. The standard deviation of a site’s annual precipitation and growing season temperatures over the past 120 years also varied dramatically (Table 1 and S1), with precipitation inter-annual standard deviation (IASD) varying from 40mm to 800m, spring temperature IASD from 0.77C to 1.95C, and summer temperature IASD from 0.35C to 1.32C (Figure 2 and S1 and Table S1). The southwest US generally had the highest precipitation IASD relative to its mean precipitation, with IASD being nearly equal to the mean precipitation in some regions (Figure 2).

**Table 1:** Summary statistics of climatic patterns relevant to the evolution of within and transgenerational plasticity. Mean (s.d.). IASD: Inter-annual standard deviation (representative of how variable conditions are between years). ACF: Autocorrelation at lags 1, 2 and, 3.

|  | Mean | IASD | ResACF-1 | ACF-1 | ACF-2 | ACF-3 |
| --- | --- | --- | --- | --- | --- | --- |
| Precipitation | 763 (443) | 145 (76) | 0.02 (0.08) | 0.04 (0.09) | 0.05 (0.13) | 0.01 (0.09) |
| Spring Temp | 10.4 (5.5) | 1.2 (0.2) | -0.04 (0.07) | -0.01 (0.08) | -0.04 (0.1) | 0.05 (0.09) |
| Summer Temp | 21.2 (4.2) | 0.9 (0.2) | 0.17 (0.1) | 0.24 (0.12) | 0.09 (0.1) | 0.11 (0.09) |

**Figure 2:**
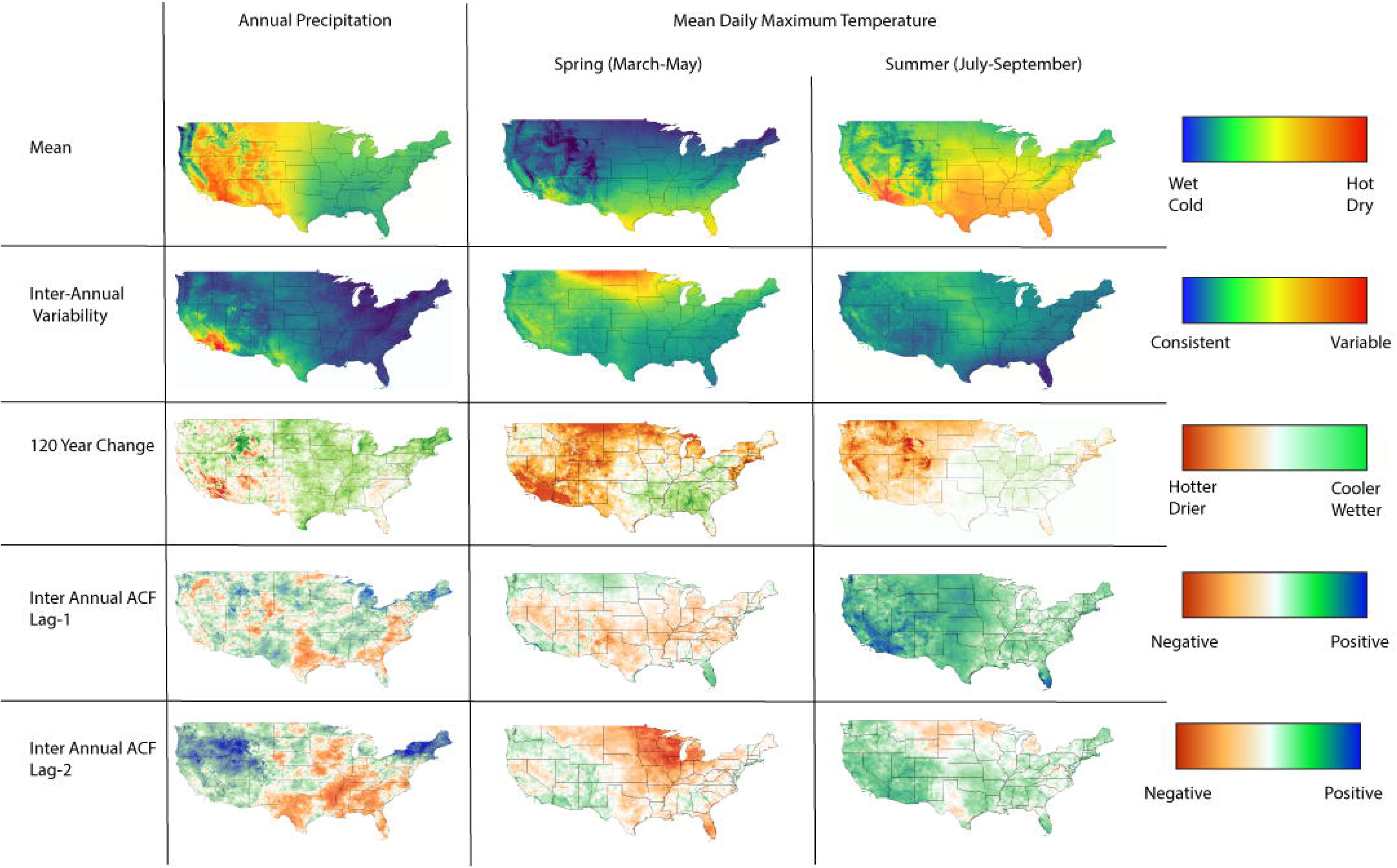
Maps depicting natural climatic variation across the conterminous US.

Directional climate change over the past 120 years was prevalent and variable across the US (Figure 2). Mean annual precipitation has declined over much of the Sierra Nevada mountain range, southern California, and other scattered regions over the last 120 years, while precipitation levels have increased in the Midwestern and much of the northeastern US. Both spring and summer temperatures have risen substantially with the exception of the southeast, where spring temperatures have decreased and summer temperatures have changed little (Figure 2). This phenomenon has been noted numerous times (Knappenberger et al. 2001; Ellenburg et al. 2016) and seems to be largely due to a switch from cropland to natural forest ecosystems across the southeastern US during the past 120 years that has led to greater transpiration cooling.

Although a variable environment is necessary for the evolution of adaptive phenotypic plasticity, it is the patterns and predictability of this variation that influence which forms of plasticity will be favored. In particular, when the grain of environmental variation is such that autocorrelations between the parental environmental cue and the offspring environment at the time of selection, mutual information will be maximized by transgenerational plasticity. We calculated autocorrelations in annual temperature and precipitation levels between successive years, which allowed us to sum across the frequencies of environmental fluctuations to capture the scale of environmental grain expected to favor transgenerational plasticity in annual species. We found that the magnitude and directions of autocorrelations on this timescale were highly variable across the US (Figure 2 and S1, Table 1 and S1).

Averaged across all sites, the precipitation autocorrelation (AC) at lag-1 (i.e., the correlation between the precipitation one year and the next) was slightly positive (mean=0.04, Table 1, Figure S1), and was reduced by half after taking linear changes in precipitation into account (mean=0.02, Table 1, Figure S1). Spatially, we found that the southeastern gulf coast was the largest region with negative lag-1 ACF (dry years tend to be followed by wet years), while the northeastern US was the largest region of substantially positive lag-1 ACF (Figure 2). Somewhat surprisingly, there were many more sites with moderately positive (62,693: lag-2 PACF > 0.2, vs. 21,671: lag-1 ACF >0.2) and negative (5,088 lag-2 PACF <-0.2 vs. 441 lag-1 ACF <-0.2) lag-2 partial autocorrelation (PAC) than lag-1 ACF. This suggest that climatic oscillations impacting annual precipitation tend to operate over more than two years in these regions, and that on a year to year basis, variation is more stochastic (leading to lower absolute lag-1 ACF).

Patterns of temperature autocorrelations extended over larger regions and were more extreme than the patterns observed for precipitation autocorrelations (Figure 2). Lag-1 ACF for spring and summer temperatures varied a great deal, with patterns of summer temperature autocorrelation substantially more positive than those of spring (summer ACF-1 mean: 0.24, spring mean: −0.01, Figure S1). In both cases, however, the western US tended to have more positive autocorrelations than the rest of the country (with the exception of southern Florida; Figure 2). The mean lag-2 PACF for spring temperature was negative (mean: −0.04, Figure 3) and more variable than lag-1 ACF (sd=0.1 vs. 0.08), with much of the north-central US displaying lag-2 PACF of less than −0.2 (Figure 2). The mean lag-2 PACF for summer temperatures was positive (mean: 0.09), but substantially lower than the mean lag-1 ACF (mean: 0.24).

**Figure 3:**
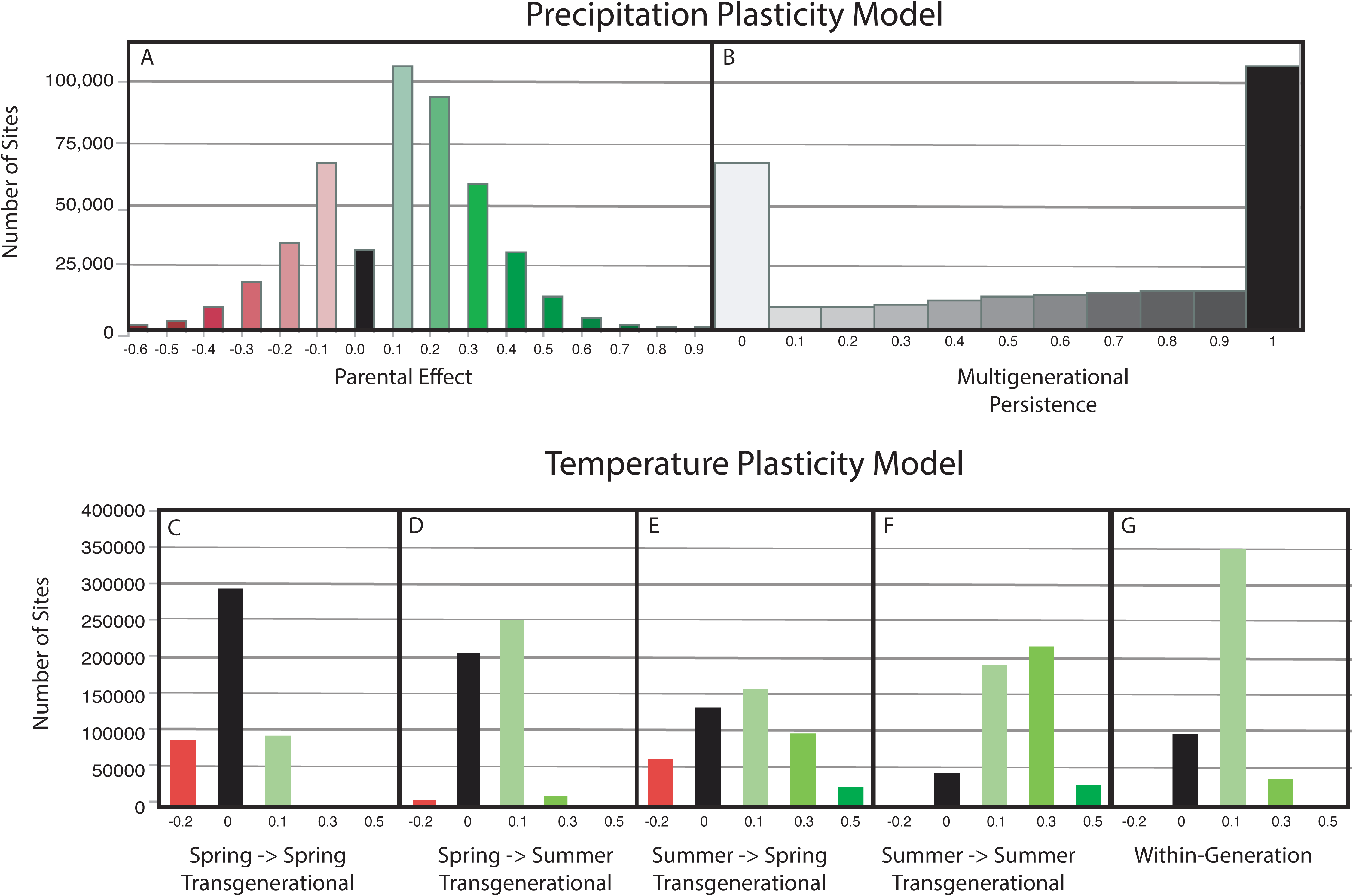
Distributions of optimal plasticity (A and B) and temperature (C-G) values across all 4km x 4km sites in the US. Histograms of optimal (A) precipitation transgenerational plasticity value (*T*) (early season) transgenerational plasticity (***m*_*EE*_**), (D) temperature spring (early season) -> summer (B) precipitation multi-generation persistence (*G*), (C) temperature spring (early season) -> spring season) transgenerational plasticity (***m*_*LE*_**), (F) temperature summer (late season) -> summer (late season) transgenerational plasticity (***m*_*EL*_**), (E) temperature summer (late season) -> spring(early season) transgenerational plasticity (***m*_*LL*_**), and within generation temperature plasticity (*W*)

Modelling work on transgenerational plasticity has often focused on positive lag-1 autocorrelations and found them to be highly correlated with optimal transgenerational plasticity. In the particular case of anticipatory transgenerational effects, the autocorrelation between the environment the parent experiences and the offspring selective environment was found to be almost perfectly tied to the evolved mean maternal effects after 50,000 simulated generations (Kuipjer et al., 2014), with similar results in a number of other studies (Tufto 2015; English et al. 2015; McNamara et al 2016). Partial-autocorrelations at lag-2 represent the additional mutual information captured by the grandparental environment, and are therefore is expected to influence the evolution of transgenerational effects that are transmitted over two generations. From these prior modeling results it is reasonable to expect that locations with high positive autocorrelations may be favorable for the evolution of transgenerational plasticity. Within these sites, areas with high lag-2 partial autocorrelations may favor the transmission of environmental information across two generations.

## Modeling Results

### Optimal levels of transgenerational plasticity: precipitation

As expected, the dramatic variability of precipitation autocorrelations across the US leads to a great deal of variation in the optimal levels of plasticity in our evolutionary models (Figure 3, Table S1). In the raw variant of the model, optimal parental effect values were positive in 314,118 cases (65%), zero in 32,352 (7%), and negative in 135,161 (28%), compared to 55%, 7%, and 38% respectively in the residual variant. The most common “parental effect” value (***m***, see Appendix 1) in the precipitation model was 0.1 (22.5% and 21.8% of sites in the raw and residual models, respectively, Figure 3a.). This level of parental effect indicates that 90% of phenotypic variance is dictated by the long-term average (genetic effects), and 10% by the difference between the parental environment and the long-term average environment. The second most common optimal value of ***m*** was 0.2 (19.15% in the raw model, 18.6% in residual model), followed by −0.1 (14.3% in the raw model, 17.2% in the residual, Figure 3a).

The multigenerational persistence (***g***, Appendix 1) of transgenerational effects was also found to vary greatly across the US with the two most common values being 1 (40.7% raw, 39.5% residual) and 0 (18.6% raw, 16.8% residual) (Figure 3b). Here, a value of 1 indicates that the precipitation one, two, and three years prior all contribute equally to the expected precipitation at a given site. A ***g*** value of 0 indicates that only the previous year’s precipitation is predictive of the current precipitation level. The remaining 40.7% of sites (in the raw variant) have intermediate optimal values of ***g***, suggesting that in these locations the precipitation of each of the past three years is informative, but information from the immediately preceding year is of the highest value (Figure 4). Interestingly, full multigenerational persistence (***g*** =1) was more frequently optimal at sites with negative transgenerational effect values compared to those with positive values (44.9% vs. 38.9%, respectively), where intermediate multigenerational persistence was more common (Figure S2).

**Figure 4:**
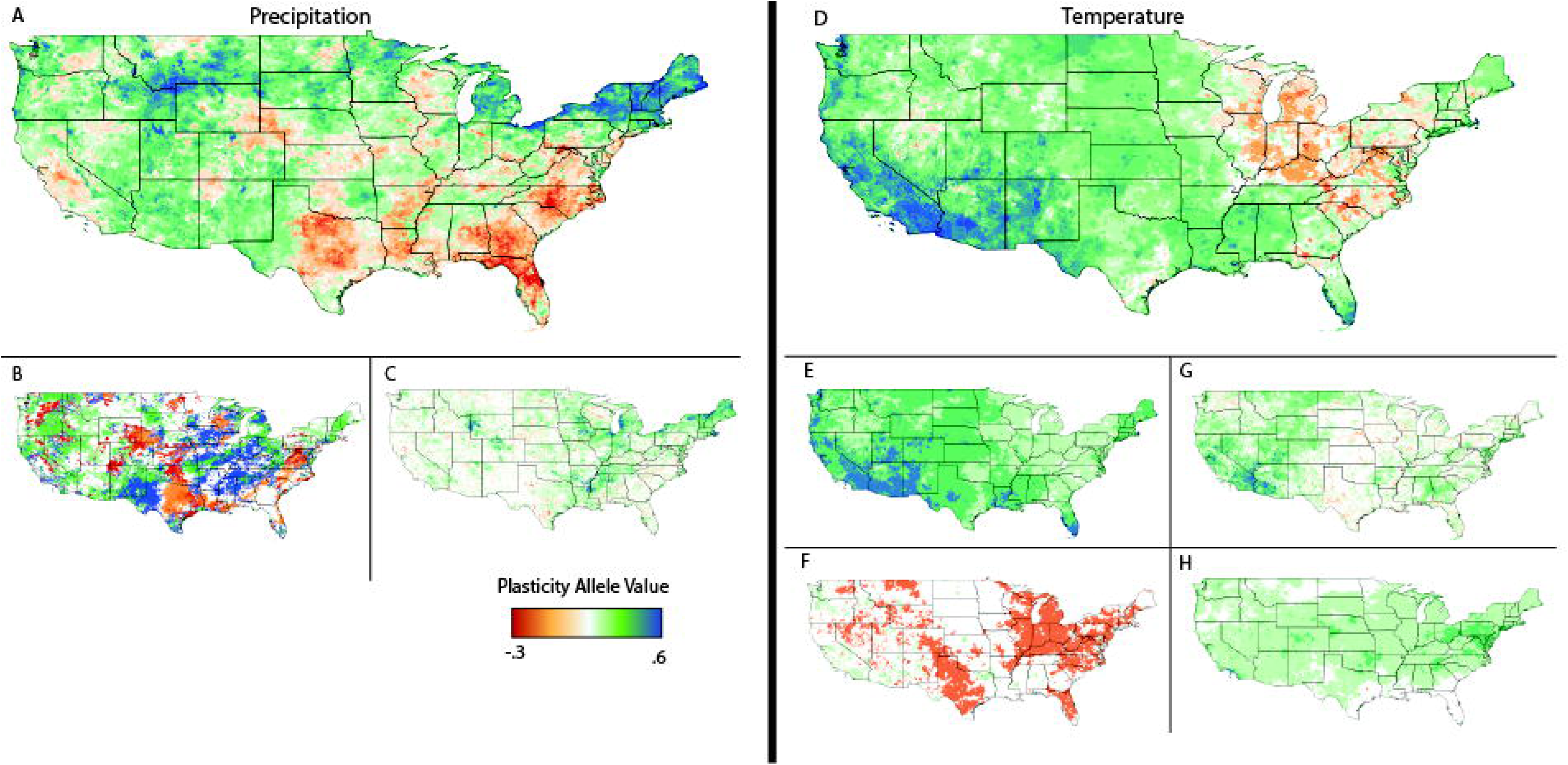
Maps coded to show patterns of variability for optimal precipitation (A-C) and temperature (D-H) plasticity values across the US. (A) Optimal transgenerational plasticity values for the one-generation transmission of precipitation level information. (B) Optimal grandparental transgenerational plasticity values coded blue (green) or red (orange) based on the direction of effect (positive or negative). White regions have an optimal multi-generation persistence (G) of 0, while red and blue both have optimal multigeneration persistence of 1, intermediate values (0>G>1) in orange and green. (C) The difference between optimal transgenerational plasticity values in the raw vs. residual variant of the mode. Higher values suggest that the primary value associated with transgenerational plasticity over the past 120 years has been associated with allowing individuals to keep up with linearly changing precipitation patterns. (D) Optimal total levels of transgenerational temperature plasticity ((M_EE_ + M_EL_ + M_LE_ + M_LL_)/2). (E) Optimal transgenerational plasticity of most extreme positive transgenerational plasticity allele. (F) Optimal transgenerational plasticity of lowest transgenerational plasticity allele. Regions in orange have at least one form of transgenerational plasticity for which negative transgenerational effects increase fitness. (G) The difference between optimal transgenerational plasticity values in the raw vs. residual variant of the mode. Higher values suggest that the primary value associated with transgenerational plasticity over the past 120 years has been associated with allowing individuals to keep up with increasing temperature. (H) Optimal within generation plasticity (W) values.

Spatial variation for optimal precipitation plasticity values largely paralleled the spatial distribution of inter-annual precipitation autocorrelation patterns (compare Figure 2a to Figure 4a). This agrees with previous modeling results that have linked autocorrelation levels with the optimal levels of transgenerational plasticity (McNamaraa et al., 2016). At the broadest level, the northern latitudes show the highest optimal transgenerational precipitation plasticity values (Figure 4a), but not necessarily multi-generation persistence of transgenerational effects (Figure 4b). Optimal transgenerational plasticity values were on average 0.057 lower in the residual variant of the model compared to the raw variant, with the vast majority of sites having equal values (56%), decreasing by 0.1 (24%), decreasing by 0.2 (8.5%), or increasing by 0.1 (4.3%). The northeastern US and the Yellowstone National Park region, where precipitation increased most (Figure 2), also saw the greatest proportion of their optimal transgenerational plasticity values diminished after factoring out linear climate change (Figure 4c). Therefore, although transgenerational plasticity has been optimal over the past 120 years in these regions, these benefits appear to be contingent upon recent warming trends.

### Optimal levels of transgenerational plasticity: Temperature

Purely positive transgenerational effects **(*m*_*EE*_ ≥ 0, *m*_*EL*_ ≥ 0, *m*_*LL*_ ≥ 0, *m*_*LE*_ ≥ 0)** of temperature were optimal in 70.2% of sites (338,327 out of 481,631) in the raw version of the model and 55.7% of sites (268,307) in the residual variant. Conversely, only 1.4% of sites (7,018) in the raw model and 3.1% (14,777) in the residual version included only negative transgenerational plasticity values. Only 0.4% (raw model) or 0.9% (residual model) of sites totally lacked transgenerational plasticity (either positive or negative) as part of the optimal strategy. The optimal strategies in the remaining sites (28% raw model, 40% residual model) comprised a mixture of positive and negative transgenerational plasticity values. Positive within-generation plasticity was favored in 79.7% of sites (383,667), compared to only 0.03% of sites (157) in which negative within-generation plasticity (***w***, Appendix 1) was favored, and 20.3% of sites (97,807) in which no within-generation plasticity was favored (Figure 3g). Optimal levels of within-generation plasticity were generally positive and minor across the US; 72% (346,603/481,631) of sites had an optimal ***w*** value of 0.1 (Table S1).

The most common optimal form of transgenerational plasticity to temperature in both the raw and residual models was the effect of late growing season temperature on the next generation’s late growing season phenotype (***m*_*LL*_**, Figure 3f, Table 2a). Effects of late season temperature on the next generation’s early season phenotype (***m*_*LE*_**) were the most variable, with a substantial number of sites having negative transgenerational plasticity values (63,881 raw, 100,267 residual) and many others having moderate (98,524 raw, 66,625 residual) and major (26,967 raw, 10,080 residual) positive values (Figure 3e). When considering the combined plasticity value profile of a site, the most common combination of plasticity values is, EE: none (0), EL: minor (0.1), LE: minor (0.1), LL: moderate (0.3), WP: minor (0.1) (Table 2b). Summing the four transgenerational plasticity alleles together we find the southwest US has the highest optimal values of transgenerational plasticity, while the Great Lakes region has the lowest optimal values (Figure 4d). In the southwestern US, where temperature increased the most over the past 120 years (Figure 2), the difference between the raw and residual model was the greatest (Figure 4g).

**Table 2:** Most common, second most common, and mean optimal plasticity values across all sites in the US for precipitation and temperature models.

|  |  | Raw |  | Residual |  |
| --- | --- | --- | --- | --- | --- |
| Climate Term |  | #1 / #2 | Mean (s.d.) | #1 / #2 | Mean (s.d.)_ |
| Precipitation | $M$ | 0.1/0.2 | 0.094 (0.22) | 0.1/0.2 | 0.036 (0.21) |
| | $G$ | 1/0 | 0.63 (0.40) | 1/0 | 0.64 (0.40) |
| Temperature | $m_{EE}$ | 0 / 0.1 | -0.016 (0.098) | 0 / -0.2 | -0.04 (0.11) |
| | $m_{EL}$ | 0.1 / 0 | 0.057 (0.073) | 0.1 / 0 | 0.045 (0.073) |
| | $m_{LE}$ | 0.1 / 0 | 0.096 (0.178) | 0 / 0.1 | 0.042 (0.17) |
| | $m_{LL}$ | 0.3 / 0.1 | 0.204 (0.133) | 0.1 / 0.3 | 0.148 (0.117) |
| | $w$ | 0.1 / 0 | 0.095 (0.072) | 0.1 / 0 | 0.092 (0.072) |

Variation in different classes of temperature autocorrelations between seasons explains a large portion of the variation in the optimal transgenerational response to temperature at a given site. For example, the autocorrelation between early season growing temperature and the next year’s late season growing temperature is the factor that explains the largest amount of variation in optimal levels of ***m*_*EL*_** (Table 3). We assessed potential tradeoffs between different forms of transgenerational plasticity to temperature by first calculating the residuals of a particular plasticity term after accounting for the effects of environmental autocorrelations, then testing the effect of the other four plasticity terms on these residuals. There was a highly significant negative association between ***m*_*LE*_** and ***m*_*EE*_** plasticity, and between ***m*_*LL*_** and ***m*_*EL*_** plasticity (Figure S3a). As higher levels of LL transgenerational plasticity were favored, the optimal levels of EL plasticity also decreased across all environmental autocorrelation values. These associations suggest that, for a given life history stage in this model, there are tradeoffs between using transgenerational information from the previous generation’s early vs. late season temperature (Table S2). For example, there are many sites where no plasticity, ***m*_*EE*_** plasticity, and ***m*_*LE*_** plasticity all have higher fitness than individuals exhibiting both *mEE* and ***m*_*LE*_** plasticity (Figure S3b).

**Table 3:** Correlations between the temperature inter-annual autocorrelations and the optimal transgenerational plasticity values.

| | $m_{EE}$ | $m_{EL}$ | $m_{LE}$ | $m_{LL}$ |
| --- | --- | --- | --- | --- |
| EE ACF | 0.757139 | -0.131583 | -0.049501 | -0.156768 |
| EL ACF | -0.036229 | 0.5968488 | -0.069245 | -0.153818 |
| LE ACF | 0.1256068 | -0.036048 | 1.3390581 | -0.067451 |
| LL ACF | 0.0135772 | -0.017808 | -0.120773 | 0.9757164 |
| Within ACF | -0.067861 | -0.160476 | 0.0595802 | 0.0701295 |

### Fitness Landscapes

In the previous analyses we used restricted parameter space to identify optimal site-specific combinations of plasticity values across the entire contiguous U.S., but further insight can be gained by comparing fitness landscapes across the full parameter space at individual sites. As both the magnitude of transgenerational plasticity and the persistence of these effects through time have been found to vary, these two parameters represent two biologically realistic components of transgenerational plasticity variation. Using the precipitation model we found that, among sites where fitness optima are located near zero transgenerational effects, a vertical fitness ridge formed that was centered near parental effect values of zero. This result is due to transgenerational persistence levels (y-axis) having a minimal impact on phenotype when parental effects are marginal. As absolute optimal parental effect values increased, however, the fitness landscape shifted from a ridge to a peak, with certain values of transgenerational persistence imparting extreme fitness advantages over others (Figure 5). Site B (North Central Minnesota) exemplifies a unique and unexpected outcome of this model: under certain conditions, there can be multiple local fitness maxima with divergent levels of transgenerational plasticity (Figure 5). Two fitness maxima exist at this site, one in which the optimal strategy comprises slightly negative parental effect values with no multigenerational persistence, and a second in which the optimal strategy comprises slightly to moderately positive parental effect values with high levels of multigenerational persistence. This situation occurs when two conditions hold: the lag-1 autocorrelation is in a different direction than the average of the lag-2 and lag-3 autocorrelations, and the absolute value of the lag-1 autocorrelation is less than the average of the lag-2 and lag-3 autocorrelations. This scenario occurs in approximately 90k out of the 480k sites, but only in 30k sites are lag-2 and lag-3 average values greater than 0.1 and therefore likely to show up as bimodal peaks in our model.

**Figure 5:**
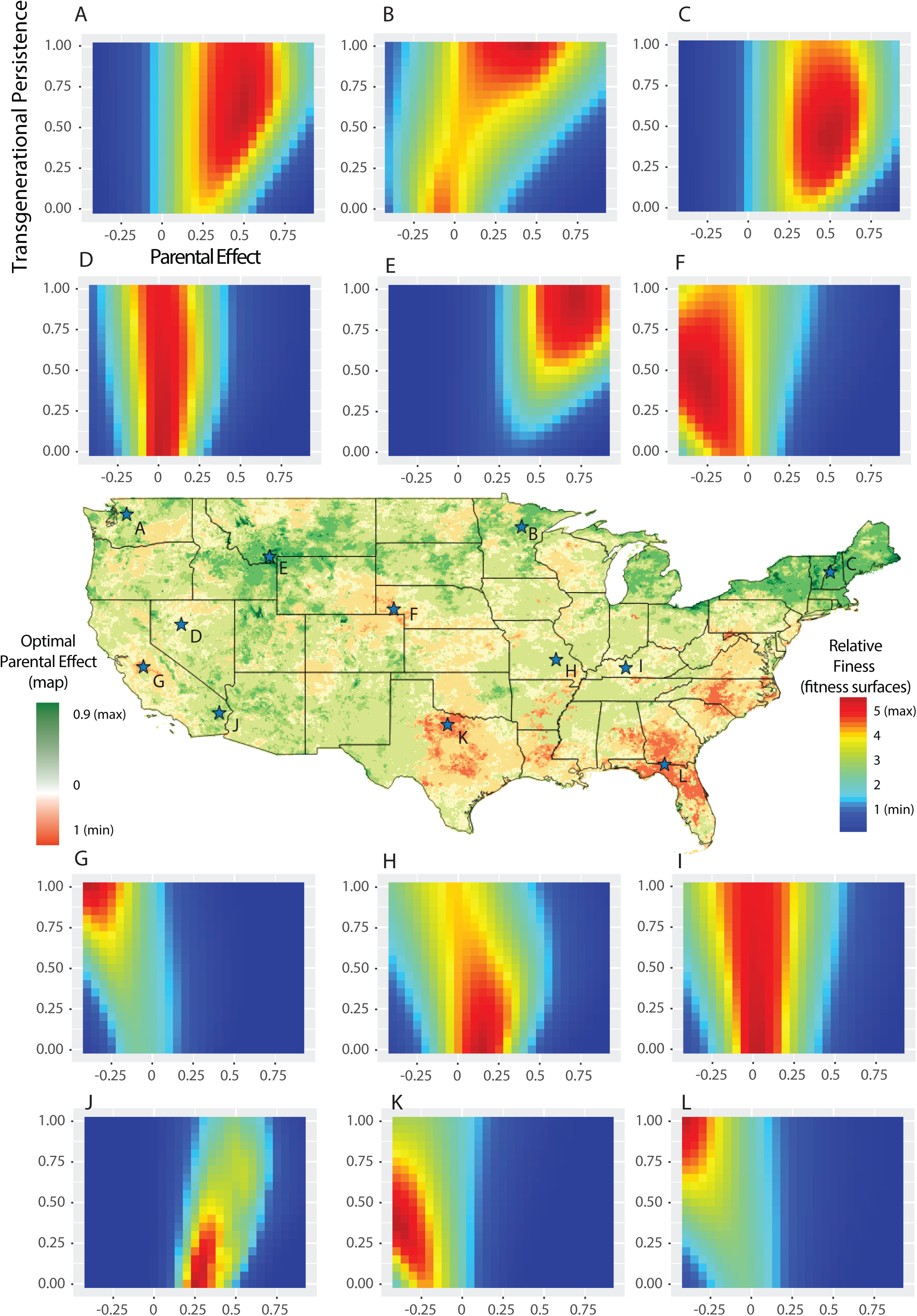
Fitness landscapes of transgenerational precipitation alleles for twelve sites across the US. Sites with low optimal parental effects (D and I) have only very subtle fitness differences associated with chanes in the multigeneration persistence (Y-axis) due to the minor role in any form of transgenerational effect on fitness in these cases. More defined fitness peaks tend to occur in areas where more substantial transgenerational effects are optimal (E, G, J, K, L). In some rare cases, bimodal fitness landscapes arise (B) where lines with either positive (with high persistence) or negative (with low persistence) transgenerational persistence have higher fitness than lines with no transgenerational inheritance.

## Discussion

Although transgenerational environmental effects on phenotypic expression have been recognized for decades (Falconer 1981; Roach and Wulff 1987), interest in these effects has surged recently due to increased appreciation for the potential role of transgenerational plasticity in adaptation (Donelson et al. 2018). Despite this renewed interest, a critical question has remained unanswered: do natural patterns of environmental variation contain fluctuations of intermediate environmental grain that favor the evolution of adaptive transgenerational plasticity? Our analysis of 120 years of climatic data from the continental U.S. revealed that such patterns are indeed widespread. Specifically, we analyzed how inter-annual variation in precipitation and temperature impacts the optimal mode of adaptation for clonally reproducing organisms with a life cycle meant to mimic that of an annual plant. When there are correlations between the parental and offspring environments, neither traditional genetic selection nor within-generation plasticity take full advantage of the available information inherent in the environment. Instead, under such correlations selection should favor the genetic evolution of mechanisms that transmit plastic responses from one generation to the next. Absent such correlations, the information provided by the parental environment may not be relevant to offspring, and indeed may prove to be maladaptive (reviewed by Herman et al. 2014).

Our modeling results revealed that the vast majority of sites in the contiguous US experienced autocorrelations in precipitation and temperature that should favor the evolution of adaptive transgenerational plasticity. As predicted by other models, the predictability of an environmental variable as measured by its autocorrelation is a major factor driving the optimal level of plasticity (e.g., Groot et al. 2017; English et al. 2015; Sultan and Spencer 2002; Scheiner 2016). Furthermore, we find that the strength and direction of autocorrelations in precipitation and temperature varied substantially across the U.S., and consequently, the optimal levels of plasticity were also highly variable. These results provide novel insight into where transgenerational effects are likely to evolve.

The environmental autocorrelation between successive generations reduces the spectra of environmental oscillations, or the grain of environmental variation, to a metric that is highly relevant to transgenerational plasticity. While the precise relationship between the level of autocorrelation and the optimal degree of transgenerational plasticity can vary depending on the precise modeling conditions, autocorrelations between parental environments and offspring selective environments are consistently associated with environments that select for transgenerational plasticity. A recent synthesis of transgenerational plasticity studies highlighted the importance of considering environmental predictability when designing experiments that test for the presence of adaptive transgenerational plasticity (Yin et al. 2019). Indeed, some experiments that failed to find evidence of adaptive transgenerational plasticity were in systems where models would not expect such effects to evolve. Our results provide a starting point for biologists looking to design experiments on natural variation in transgenerational plasticity.

### Precipitation

Local adaptation to variable water regimes has been a major focus of plant evolutionary ecology for many years. This literature shows that plants have evolved a wide range of physiological, phenological, and morphological adaptations to handle site-specific patterns of water availability (Kooyers 2015). These adaptive phenotypes may be expressed constitutively or may be induced by an environmental cue that predicts a change in water availability later in the life of the organism. Increasingly, experimental studies show that the parental soil moisture regime can also adaptively influence the development of progeny (e.g., Alsdurf et al. 2013; Alsdurf et al. 2015), providing a third route by which plants can fine tune the phenotypes of their offspring to local soil-moisture levels. For instance, in Massachusetts genotypes of the annual plant *Polygonum persicaria*, offspring of drought-stressed parents make more extensive root systems and deploy them faster in response to drought as compared to offspring of well-watered parents. This drought-induced change in growth and development can be inherited for at least two generations, resulting in increased survival of grand-offspring under severe drought stress (Sultan et al 2009; Herman et al. 2012). Furthermore, these epigenetic effects of drought are genetically variable in *P. persicaria*: some genotypes strongly increase root length and biomass in response to parental drought, while other genotypes do so only moderately or not at all (Herman and Sultan 2016).

Our analysis revealed substantial and spatially variable interannual autocorrelations in precipitation, indicating that precipitation levels in one year are often predictive of precipitation levels up to three years later. For example, across the coterminous U.S., lag-1 interannual precipitation autocorrelations varied from moderately negative (−0.27) to strongly positive (0.69), including some values near zero. In turn, the optimal direction and strength of transgenerational effects of precipitation also varied. Positive parental effects, wherein individuals are developmentally predisposed to perform better in environments that match their parents’ environment, were optimal across more than twice as many regions (65% of sites) as negative transgenerational effects (28% of sites), wherein individuals perform better in a different environment than their parents. Relatively strong parental effect values of 0.3 or higher were optimal in nearly 30% of sites. By contrast, complete absence of parental effects was favored in only 7% of sites.

Multigenerational persistence values of 0 (18.7% of sites) and 1 (40.7% of sites) were most common, representing strategies in which transgenerational effects lasted only a single generation or persisted fully to the third generation, respectively. The remaining persistence values were somewhat evenly distributed between 0 and 1 and represent strategies in which environmental information gets passed through three generations, but the environment of recent years is weighted more heavily.

The optimal level of transgenerational effects varied on multiple scales. On the largest scale, we found that the western and northern US experience conditions that select for the highest levels of transgenerational plasticity (Figure 4a). There was a striking contrast between the northeast, where positive transgenerational plasticity was generally optimal, and the southeast, where negative transgenerational plasticity predominated. On these intermediate to large spatial scales, it is likely that natural selection could counteract the homogenizing force of gene flow to generate patterns of locally adaptive transgenerational plasticity to precipitation. Experiments designed to compare transgenerational plasticity to precipitation in individuals derived from these north/south or east/west clines would provide novel evidence for climatic patterns shaping the system of inheritance in individuals. There was also considerable variation in optimal levels of transgenerational plasticity on much finer scales. In some cases, levels of transgenerational plasticity were highly divergent between adjacent sites (e.g., in Texas and Minnesota). In these cases, and particularly for outcrossing species, it is less likely that natural selection would be able to counteract gene flow, perhaps limiting the locally adaptive evolution of transgenerational plasticity.

### Temperature

Temperature is vitally important to plant function and fitness, as it impacts the rate of physiological reactions, cues developmental transitions, and in extremes can cause stress and mortality. Plants adapt to variable temperature regimes in a host of ways, including the production of heat shock proteins and cold-response factors, and the development of morphologies that mitigate the experience of temperature extremes. Experimental studies have identified adaptive plastic responses to temperature changes, both within and across generations. For example, ambient temperature in *Arabidopsis thaliana* has been shown to influence the expression and splicing of hundreds of genes, leading to changes in histone methylation (Pajoro et al. 2017), and shifts in flowering time (Donohue 2009) and other phenotypes (Adams et al. 2016) in genotype-specific ways. Additionally, recent work has demonstrated that effects of temperature on *A. thaliana* plants persist for multiple generations (Whittle et al. 2009; Suter and Widmer 2013a; Suter and Widmer 2013b; Groot et al. 2017). In order for these responses to adaptively match phenotypes with environments, there must be substantial correlations in temperature within and between growing seasons.

We found significant autocorrelations in temperature, both within and between years. Within a single growing season, temperatures early and late in the growing season tended to be positively correlated across the U.S. Furthermore, we found that the temperatures of the late growing season months (July, August, September) were generally strongly autocorrelated between successive years. Interannual correlations between the temperatures of the early growing season months (February, March, April) were often much lower. As expected, we find that, at a given site, the strength of the correlation between the average temperature during the season in which information is gathered and the average temperature during the season when selection occurs is highly predictive of both the type and degree of plasticity that will be favored. For example, warmer than average springs were very often followed by hotter than average summers, and this information yielded benefits via within-generation responses to temperature in many sites. The optimal strategy in more than 99% of sites across the U.S. contained some form of transgenerational plasticity, suggesting that environmental oscillations provide valuable information that allows transgenerational plasticity to improve the match between phenotypes and temperature regimes.

The most common form of transgenerational plasticity in this model was late-growing season temperature impacting the following generation’s phenotype late in the growing season, which matches our expectations based on the patterns of temperature autocorrelation. Interestingly, patterns of environmental oscillations lead to favorable strategies in which the current late-season phenotype was more strongly impacted by the previous late-season temperature than it was by the current generation early season temperature. Indeed, this pattern was found in over half of the regions considered (270k/480k). Although intuition suggests that more recent information is of higher value, this result suggests that parental environments can be more predictive of offspring selective environments than environmental cues early in the offspring generation. This result stems from the cyclic nature of seasonal environments (Auge et al. 2017). Since autocorrelations between consecutive early growing seasons were generally low, it is not surprising that effects of early growing season temperatures on phenotypes in the following early growing season was the least common form of plasticity and was in the negative direction more often than the positive. Other forms of transgenerational plasticity were present at intermediate levels and varied across the US.

The west coast of the US and southern Florida experienced the highest optimal transgenerational plasticity values. Because these regions are due east of large bodies of water, their climates are heavily influenced by maritime airflow including the prevailing westerlies, loop current, and Coriolis affect (Lorentz 1966). As water has a substantially higher heat capacity than either rock or soil, the location of these land masses downstream of maritime air may predispose them to temperature autocorrelations between years, but whether this result is universal will take studies on other continents. These areas may be primed for large-scale, community level comparisons of transgenerational plasticity. Comparisons of transgenerational plasticity in individuals found on the west vs. east coast could shed light on the generality of these patterns across a diversity of annual plants and other taxa.

We found highly variable associations between late growing season temperature and the following generation’s early growing season temperature. This result is intriguing because the temperature experienced during seed maturation strongly influences the dormancy and germination behavior of seeds, with cascading effects throughout the life cycles of annual plants (Donohue 2009; Burghardt et al. 2016). Consequently, site-specific correlations between maternal late-season temperature and the early-season temperature in the next generation may select for divergent, site-specific effects of maternal temperature on germination. Intriguingly, parental effects of temperature on germination and flowering time are highly genetically variable in *A. thaliana* (Burghardt et al. 2016; Kerdaffrec and Nordburg 2017; Groot et al. 2017). In *Plantago lanceolata*, such genotype-by-maternal temperature effects persist throughout the offspring life cycle to generate variation in reproduction in the field (Lacey and Herr 200). Our results suggest that genetic variation for maternal effects may derive in part from variable selection imposed by differences among sites in temperature correlations (see also Groot et al. 2017).

## Common Themes and Future Directions

Although our precipitation and temperature models yielded distinct insights into the dynamics of each of these factors, common themes emerged in both sets of analyses. For example, we found higher levels of inter-annual autocorrelation, and therefore more prominent transgenerational effects, at northern latitudes and along coastal regions within both models. Studies that compare patterns of transgenerational plasticity across such large geographic regions will be necessary to determine whether underlying differences in environmental patterns do in fact drive differences in transgenerational plasticity. While the scale of gene flow varies greatly among species, these large-scale patterns generate large contiguous regions with divergent optimal levels of transgenerational plasticity that should provide ample opportunity for natural selection to drive the evolution of transgenerational effects even in the face of gene flow. For example, in our temperature model, the western half of the US represents a contiguous region where positive transgenerational effects are expected to evolve, while the neighboring great lakes region is many thousands of square miles in area with negative optimal transgenerational plasticity. These large regional differences should allow selection to produce divergent transgenerational norms of reaction; future studies explicitly modeling the migration and evolutionary parameters of specific species will be necessary to test these predictions in different scenarios.

Another common finding of both the temperature and precipitation models is that transgenerational effects are expected to provide greater benefits in changing climates relative to purely oscillating climates, in which linear climate change has been removed (i.e., the residual models). These results suggest that transgenerational effects may have an important role in adaptation to human-induced climate change, and that rapid climate change should select for more transgenerationally plastic individuals. However, there is an important caveat. In our models we assume that genotypes are uniform in their mean phenotype, and do not allow for mutations that could lead to genetic adaptation to changing conditions. The potential for transgenerational plasticity to either promote or hinder genetic adaptation has been explored (Day and Bonduriansky 2011), but our models do not address this issue. In the absence of genetic evolution, it follows that if there is a linear trend towards hotter or drier years in addition to climatic oscillations (as in the raw model variants), then there is more transgenerational information relative to a situation in which only climatic oscillations are occurring (residual model variants). Theory indicates that these dynamics become much more complex when local genetic adaptation to changing conditions is allowed to occur along with plastic responses (Groot et al. 2017). For instance, in some scenarios transgenerational effects can increase fitness in the short term, while reducing it in the long term (Hoyle and Ezard 2012).

Temperature and precipitation autocorrelations likely stem in part from the same broad-scale climatic oscillations, such as the El Niño Southern Oscillation (Yang et al. 2018), the Quasi-biennial oscillation (Baldwin et al. 2001), and the Pacific Decadal Oscillation (Mantua and Hare 2002; Newman et al. 2016). Aside from these climatic oscillations, autocorrelations will arise due simply to “red” or “pink” noise in which rare, large events and common, small events have equal power in explaining variation (Szendro et al. 2001). It has been demonstrated that even without clear underlying phenomena explaining variation, pink-noise is often the model that best explains patterns of ecological and abiotic time-series variation (Halley 1996). These oscillations and general patterns of red noise will interact with each other to varying degrees across different regions of the US, leading to variable levels of autocorrelation at all lags for both precipitation and temperature.

Furthermore, because temperature and precipitation interact to alter moisture availability, it is likely that organisms do not process temperature and precipitation information independently, but rather use them in tandem along with other sources of information to fine tune phenotypes for the most likely future environment. For instance, temperature influences water availability by influencing rates of evaporation and transpiration. Interactions between temperature and water availability also shape the collection of herbivores, pathogens, and competitors present in a given locality. Understanding how these environmental factors jointly influence the expression of transgenerational plasticity is an important goal for future research.

A key element of this research direction is to study environmental (auto)correlations at fine scales in the context of dispersal distances. It is possible that transgenerational plasticity may be a more common mode of adaptation for organisms with short dispersal distances, in which parents and offspring are more likely to grow and develop in similar microsites. Finally, differences in life history strategies and generation times will alter the timescales and types of environmental autocorrelations relevant to transgenerational plasticity.

A recent meta-analysis of 1,170 transgenerational plasticity effect sizes found that there was substantial evidence for adaptive transgenerational plasticity, but that these effects varied according to the type of trait that was considered, the environmental context, and the taxonomic and life-history group of the focal organism (Yin et al., 2019). In particular, this meta-analysis found that annual plants displayed the most substantial evidence for adaptive transgenerational plasticity, and that physiological traits showed the highest evidence for adaptive plasticity to parent environments. The finding that annual plants displayed the greatest degree of transgenerational plasticity is consistent with their limited mobility and short-life cycle, both of which increase the likelihood that offspring experience similar environments to their parents. The mean effect size found in this study for annual plants was 0.163 for reproductive traits and 0.216 for physiological traits, which is consistent with our modeling results. We found that the mean inter-annual summer temperature autocorrelation was 0.24 and 0.17 before and after factoring out linear effects of climate change, with optimal transgenerational effect sizes in our temperature model ranging from 0 to 0.3. While Yin et al. 2019 did not consider differences between environmental variables, inter-annual temperature autocorrelations could drive autocorrelations in a diversity of selective pressures. Taken together with our modeling results, this meta-analysis indicates that observed strengths of transgenerational effects in annual plants are in line with the predictions made by patterns of autocorrelations observed in nature. Similarly, Yin et al. (2019) found that short-lived invertebrates were the second most likely group to express transgenerational plasticity, suggesting that the capacity to transmit epigenetic information between generations is not phylogenetically limited. For longer-lived taxa, such inter-annual autocorrelations would be relatively fine-grained, and thus more likely to select for within-generation rather than transgenerational plasticity. Future studies modeling the evolution of transgenerational plasticity in individuals with disparate life histories will be critical for better understanding the evolution of these environmental effects.

## Conclusion

In summary, we demonstrate that patterns of climatic variation in nature may favor the adaptive evolution of transgenerational plasticity in organisms with approximately annual generation times, such as annual plants. Our models indicate that differing patterns of climatic oscillations across the US lead to strikingly different optimal patterns of within- and transgenerational plasticity. Thus, for a given species, one may expect that environmental variation across its range not only selects for different locally adapted mean trait values, but also different classes and magnitudes of plasticity. Perhaps the most meaningful result of this study is that the climatic patterns across relatively small geographic regions vary so dramatically that the optimal value of transgenerational plasticity ranges from extremely high to non-existent. It should therefore be expected that although many species, environmental variables, or phenotypes of interest may show no evidence of transgenerational plasticity, such results may be due to their specific ecological situation rather than a fundamental biological limitation. This applies equally strongly to the other side of the coin: because a single population or species expresses strong transgenerational plasticity does not mean that transgenerational effects are a universally key driver of evolutionary processes. Rather, variation in transgenerational plasticity should be expected, just as genetic variation is ubiquitous in natural populations. Transgenerational plasticity is best considered in the specific ecological and evolutionary context of the study organism, and broad generalizations about the role of these effects in evolution should be avoided until considerably more field data are in hand. The results described here provide a source of testable predictions for geographical variation in this mode of adaptation.

## Acknowledgments

We thank the Blackman and Bergelson lab groups for comments on earlier versions of the manuscripts. We also thank the UC Berkeley BRC computing core faculty for computing resources and aid with running simulations.

## Data Accessibility

All modeling inputs and results are publicly available within the supplemental tables and code on github.

---

Figure S1: Distribution of climatic summary statistics across the 481k 4×4km grids in the US. Dotted lines at 0 for autocorrelation histograms.

Figure S2: Mosaic plot showing the frequency of specific combinations of optimal transgenerational effects and multi-generational persistence values. More subtle transgenerational effects more frequently only have a single generation of persistence (G=0), while more extreme transgenerational effects tend to coincide with full transgenerational persistence (G=1) where each of the prior three years contribute equally. Additionally, positive transgenerational effects were more likely to have intermediate levels of persistence than negative transgenerational effects.

Figure S3: Figures demonstrating tradeoffs between classes of plasticity. (A) Tradeoffs between early-late (m12) and late-late (m22) transgenerational plasticity in relation to the autocorrelations in summer temperature. Generally, as inter-annual summer autocorrelations increase, so too does the frequency of minor and moderate early-late transgenerational plasticity. However, we also find that areas that favor higher levels of late-late transgenerational plasticity tend to favor lower levels of early-late plasticity compared to other sites with similar levels of temperature autocorrelation. (B) Within a single site there are many examples of localities where either early-late or late-late transgenerational plasticity lead to fitness increases relative to clones with no plasticity, but individuals expressing both forms of plasticity have the lowest fitness of all.

## References

1. Adams, W.W., J.J. Stewart, C.M. Cohu, O. Muller, & B. Demmig-Adams. 2016. Habitat temperature and precipitation of Arabidopsis thaliana ecotypes determine the response of foliar vasculature, photosynthesis, and transpiration to growth temperature. Front. Plant Sci. 7:1026. doi. 10.3389/fpls.2016.01026.

2. Agrawal, A., C. Laforsch, and R. Tollrian. 1999. Transgenerational induction of defences in animals and plants. Nature 401:60–63.

3. Akkerman, K., A. Sattirin, J. K. Kelly, and A. G. Scoville. 2016. Transgenerational plasticity is sex-dependent and persistent in yellow monkeyflower (Mimulus guttatus). Environmental Epigenetics. doi: 10.1093/eep/dvw003

4. Alsdurf, J. C. Anderson, D.H. Siemens. 2015. Epigenetics of drought-induced trans-generational plasticity: consequences for range limit development. AoB Plants 8:plv146. doi: 10.1093/aobpla/plv146.

5. Alsdurf, J. D., T. J. Ripley, S. L. Matzner, and D. H. Siemens. 2013. Drought-induced trans-generational tradeoff between stress tolerance and defence: consequences for range limits? AoB Plants 5:plt038.

6. Auge, G. A., L. D. Leverett, B. R. Edwards, and K. Donohue. 2017. Adjusting phenotypes via within- and across-generational plasticity. New Phytol. 216:343–349.

7. Baldwin, M.P., L.J. Gray, T.J. Dunkerton, K. Hamilton, P.H. Haynes, W.J. Randel, J.R. Holton, M.J. Alexander, I. Hitora, T. Horinouchi, D.B.A. Jones, J.S. Kinnersley, C. Marquardt, K. Sato, M. Takahashi. 2001. The quasi-biennial oscillation. Rev. Geophys. 39:179–229. doi: 10.1029/1999rg000073.

8. Bilichak, A., Y. Ilnystkyy, J. Hollunder, and I. Kovalchuk. 2012. The Progeny of Arabidopsis thaliana Plants Exposed to Salt Exhibit Changes in DNA Methylation, Histone Modifications and Gene Expression. PLoS ONE 7:e30515. doi:10.1371/journal.pone.0030515.

9. Bonduriansky, R. and T. Day. 2009. Nongenetic inheritance and its evolutionary implications. Annu. Rev. Ecol., Evol. Syst. 40:103–125.

10. Burghardt, L.T., B.R. Edwards, K. Donohue. 2016. Multiple paths to similar germination behavior in Arabidopsis thaliana. New Phyt. 209:1301–1312. doi: 10.1111/nph.13685.

11. Case, A. L., E. P. Lacey, and R. G. Hopkins. 1996. Parental effects in *Plantago lanceolata* L. II. Manipulation of grandparental temperature and parental flowering time. Heredity 76:287–295.

12. Chen, M., MacGregor, D.R., Dave, A., Florance, H., Moore, K., Paszkiewicz, K., Smirnoff, N., Graham, I.A. and Penfield, S., 2014. Maternal temperature history activates Flowering Locus T in fruits to control progeny dormancy according to time of year. Proceedings of the National Academy of Sciences. 111(52), pp.18787–18792.

13. Colicchio, J. 2017. Transgenerational effects alter plant defence and resistance in nature. J. Evol. Biol. 30:664–680. doi: 10.1111/jeb.13042.

14. Colicchio, J., F. Miura, J. Kelly, T. Ito, L. Hileman. DNA methylation and gene expression in Mimulus guttatus. 2015. BMC Genomics 16:507. doi: 10.1186/s12864-015-1668-0.

15. Conrath, U., G. J. Beckers, C. J. Langenbach, and M. R. Jaskiewicz. 2015. Priming for enhanced defense. Annu Rev Phytopathol 53:97–119.

16. Daly, C., W. P. Gibson, G. H. Taylor, G. L. Johnson, and P. Pasteris. 2002. A knowledge-based approach to the statistical mapping of climate. Climate Research 22:99–113.

17. Dantzer, B., B. Dantzer, A. E. M. Newman, A. E. M. Newman, R. Boonstra, R. Boonstra, R. Palme, R. Palme, S. Boutin, S. Boutin, M. M. Humphries, M. M. Humphries, A. G. McAdam, and A. G. McAdam. 2013. Density Triggers Maternal Hormones That Increase Adaptive Offspring Growth in a Wild Mammal. Science. 340:1215–1217. doi: 10.1126/science.1235765.

18. Day & Bonduriansky. 2011. A unified approach to the evolutionary consequences of genetic and non-genetic inheritance. Am Nat. 178:E18–E36. doi: 10.1086/660911.

19. Dey, S., x, S.R. and Teotonio, H., 2016. Adaptation to temporally fluctuating environments by the evolution of maternal effects. PLoS biology, 14(2), p.e1002388.

20. Donaldson-Matasci, M., C. Bergstrom, and M. Lachman. 2013. When unreliable cues are good enough. Am. Nat. 182:313–327.

21. Donelson, J.M., S. Salinas, P.L. Munday, L.N.S. Shama. 2018. Transgenerational plasticity and climate change experiments: where do we go from here? Glob. Change. Biol. 24:13–34. doi: 10.1111/gcb.13903.

22. Donohue, K. 2009. Completing the cycle: maternal effects as the missing link in plant life histories. Phil. Trans. R. Soc. B. 364:1059–1074. doi: 10.1098/rstb.2008.0291.

23. Ellenburg, W.L., R.T. McNider, J.F. Cruise, J.R. Christy. 2016. Towards an understanding of the Twentieth-Century cooling trend in the southeastern United States: biogeophysical impacts of land-use change. Earth Interactions 20. doi: 10.1175/ei-d-15-0038.1.

24. English, S., I. Pen, N. Shea, and T. Uller. 2015. The information value of non-genetic inheritance in plants and animals. PLoS ONE 10:e0116996. doi:10.1371/journal.pone.0116996.

25. Falconer, D. S. 1981. Introduction to Quantitative Genetics. London: Longman.

26. Fisher, R.A. 1930. The Genetical Theory of Natural Selection. Clarendon, Oxford.

27. Furrow, R. and M. Feldman. 2014. Genetic variation and the evolution of epigenetic regulation. Evolution 68:673–683.

28. Galloway, L., J. Etterson, & McGlothlin. 2009. Contribution of direct and maternal genetic effects to life-history evolution. New Phytol. 183:826–838. doi: 10.1111/j.1469-8137.2009.02939.x

29. Ganguly, D.R., P.A. Crisp, S.R. Eichten, B.J. Pogson. 2017. The Arabidopsis DNA Methylome is Stable Under Transgenerational Drought Stress. Plant Phys. 175:1893–1912. doi: 10.1104/pp.17.00744.

30. Ghalambor, C.K., J.K. McKay, S.P. Carroll, & D.N. Reznick. 2007. Adaptive versus non-adaptive phenotypic plasticity and the potential for contemporary adaptation in new environments. Funct. Ecol. 21:394–407.

31. Graham, J.K., Smith, M.L. and Simons, A.M., 2014. Experimental evolution of bet hedging under manipulated environmental uncertainty in Neurospora crassa. Proc. R. Soc. B. 281(1787): pp.20140706.

32. Greenspoon, P. and H. Spencer. 2018. The evolution of epigenetically mediated transgenerational plasticity in a subdivided population. Evolution. doi: 10.1111/evo.13619.

33. Groot, M.P., Kubisch A., Ouborg, N.J., Pagel, J., Schmid, K.J., Verger, P., Lampei, C. 2017. Transgenerational effects of mild heat in Arabidopsis thaliana show strong genotype specificity that is explained by climate at origin. New Phyt. doi: 10.1111/nph.14642.

34. Halley, J. 1996. Ecology, evolution, and 1/f noise. Trends Ecol. Evol. 11:33–37.

35. Herman, J. J. and S. E. Sultan. 2011. Adaptive transgenerational plasticity in plants: case studies, mechanisms, and implications for natural populations. Front. Plant Sci. 2:1–10. doi: 10.3389/fpls.2011.00102.

36. Herman, J. J. and S. E. Sultan. 2016. DNA methylation mediates genetic variation for adaptive transgenerational plasticity. Proc. R. Soc B. 283:20160988. doi: 10.1098/rspb.2016.0988.

37. Herman, J. J., S. E. Sultan, T. Horgan-Kobelski, and C. Riggs. 2012. Adaptive transgenerational plasticity in an annual plant: Grandparental and parental drought stress enhance performance of seedlings in dry soil. Integr. Comp. Biol. 52:77–88.

38. Herman, J.J., H.G. Spencer, K. Donohue, & S.E. Sultan. 2014. How stable ‘should’ epigenetic modifications be? Insights from adaptive plasticity and bet hedging. Evolution 68:632–643. doi: 10.1111/evo.12324.

39. Holeski, L. M., G. Jander, and A. A. Agrawal. 2012. Transgenerational defense induction and epigenetic inheritance in plants. Trends Ecol. Evol. 27:618–626.

40. Holeski, L.M. 2007. Within and between generation phenotypic plasticity in trichome density of Mimulus guttatus. J. Evol. Biol. 20:2092–2100. doi: 10.1111/j.1420-9101.2007.01434.x.

41. Houri-Zeevi, L. and Rechavi, O., 2017. A matter of time: small RNAs regulate the duration of epigenetic inheritance. Trends in Genetics. 33(1), pp.46–57.

42. Hoyle, R. B. and T. H. Ezard. 2012. The benefits of maternal effects in novel and in stable environments. J R Soc Interface 9:2403–2413.

43. Jablonka, E. 2013. Epigenetic inheritance and plasticity: The responsive germline. Prog. Biophys. Mol. Biol. 111:99–107.

44. Kerdaffrec, E. & M. Nordborg. 2017. The maternal environment interacts with genetic variation in regulating seed dormancy in Swedish Arabidopsis thaliana. PLoS ONE 12:e0190242. doi: 10.1371/journal.pone.0190242.

45. Knappenberger, P.C., P.J. Michaels, R.E. Davis. 2001. Nature of observed temperature changes across the United States during the 20th century. Climate Res. 17:45–53.

46. Kooyers, N.J. 2015. The evolution of drought escape and avoidance in natural herbaceous populations. Plant Sci. 234:155–162. doi: 10.1016/j.plantsci.2015.02.012.

47. Kuijper, B. and R. B. Hoyle. 2015. When to rely on maternal effects and when on phenotypic plasticity? Evolution 69:950–968.

48. Kuijper, B., R. A. Johnstone, and S. Townley. 2014. The evolution of multivariate maternal effects. PLoS Comput. Biol. 10:e1003550.

49. Lacey, E. & D. Herr. 2000. Parental effects in Plantago lanceolata L. III. Measuring parental temperature effects in the field. Evolution 54:1207–1217.

50. Lachmann, M. and E. Jablonka. 1996. The inheritance of phenotypes: an adaptation to fluctuating environments. J. of Theor. Biol. 181:1–9.

51. Leimar, O. and J. M. McNamara. 2015. The evolution of transgenerational integration of information in heterogeneous environments. Am. Nat. 185:E55–E69. doi: 10.1086/679575.

52. Lopez Sanchez, A., J. H. Stassen, L. Furci, L. M. Smith, and J. Ton. 2016. The role of DNA (de)methylation in immune responsiveness of Arabidopsis. Plant J. doi: 10.1111/tpj.13252.

53. Lorentz, E. The circulation of the atmosphere. 1966. 2017Amer. Scientist 54:402–420.

54. Mantua, N. & S. Hare. 2002. The Pacific Decadal Oscillation. J. Oceanography 58:35–44.

55. Mousseau, T. A. and C. W. Fox. 1998. Maternal effects as adaptations. Oxford University Press, New York.

56. Newman, M., M.A. Alexander, T.R. Ault, K.M. Cobb, C. Deser, E. Di Lorenzo, N.J. Mantua, A.J. Miller, S. Minobe, H. Nakamura, N. Schneider, D.J. Vimont, A.S. Phillips, J.D. Scott & C.A. Smith. 2016. The Pacific Decadal Oscillation, revisited. 2016. J. Climate 29:4399–4427.

57. Nicotra, A. B., O. K. Atkin, S. P. Bonser, A. M. Davidson, E. J. Finnegan, U. Mathesius, P. Poot, M. D. Purugganan, C. L. Richards, F. Valladares, and M. v. Kleunen. 2010. Plant phenotypic plasticity in a changing climate. Trends Plant Sci. 15:684–692.

58. Pajoro, A., E. Severing, G.C. Angenent, & R.G. H. Immink. 2017. Histone H3 lysine 36 methylation affects temperature-induced alternative splicing and flowering in plants. Genome Biology. 18:102. doi: 10.1186/s13059-017-1235-x.

59. PRISM Climate Group, Oregon State University, http://prism.oregonstate.edu, created 4 Feb 2004.

60. Prizak, R., T. H. Ezard, and R. B. Hoyle. 2014. Fitness consequences of maternal and grandmaternal effects. Ecol. and Evol. 4:3139–3145.

61. Proulx, S. and H. Teotonio. 2017. What kind of maternal effects can be selected for in fluctuating environments? Am Nat. 189:E118–E137. doi: 10.1086/691423.

62. Quantum GIS geographic information system. 2012. Open source geospatial foundation project. Free Software Foundation, India.

63. Räsänen, K. and L. Kruuk. 2007. Maternal effects and evolution at ecological timescales. Funct. Ecol. 21:408–421.

64. Rasmann, S., M. De Vos, C. L. Casteel, D. Tian, R. Halitschke, J. Y. Sun, A. A. Agrawal, G. W. Felton, and G. Jander. 2012. Herbivory in the Previous Generation Primes Plants for Enhanced Insect Resistance. Plant Physiol. 158:854–863.

65. Rechavi, O., G. Minevich, and O. Hobert. 2011. Transgenerational inheritance of an acquired small RNA-based antiviral response in C. elegans. Cell 147:1248–1256.

66. Roach, D., and Wulff, R. 1987. Maternal effects in plants. Annu. Rev. Ecol. Syst. 18, 209–235.

67. Scheiner, S. 2016. Habitat Choice and Temporal Variation Alter the Balance between Adaptation by Genetic Differentiation, a Jack-of-All-Trades Strategy, and Phenotypic Plasticity. Am. Nat. 187:633–646. doi: 10.1086/685812.

68. Scheiner, S. M. 1993. Genetics and Evolution of Phenotypic Plasticity. Annu. Rev. Ecol. Syst. 24:35–68.

69. Schlichting, C. and M. Pigliucci. 1998. Phenotypic evolution: a reaction norm perspective. Sinauer, Sunderland, Massachusetts.

70. Schmitt, J., J. Niles, and R. Wulff. 1992. Norms of reaction of seed traits to maternal environments in Plantago lanceolata Am. Nat. 139:451–466.

71. Shea, N., I. Pen, and T. Uller. 2011. Three epigenetic information channels and their different roles in evolution. J. Evol. Biol. 24:1178–1187.

72. Sikkink, K.L., Ituarte, C.M., Reynolds, R.M., Cresko, W.A. and Phillips, P.C., 2014. The transgenerational effects of heat stress in the nematode Caenorhabditis remanei are negative and rapidly eliminated under direct selection for increased stress resistance in larvae. Genomics. 104(6):438–446.

73. Slaughter, A., X. Daniel, V. Flors, E. Luna, B. Hohn, and B. Mauch-Mani. 2012. Descendants of Primed Arabidopsis Plants Exhibit Resistance to Biotic Stress. Plant Physiol. 158:835–843.

74. Stjernman, M. and T. J. Little. 2011. Genetic variation for maternal effects on parasite susceptibility. J. Evol. Biol. 24:2357–2363.

75. Sultan, S. E. 1996. Phenotypic plasticity for offspring traits in Polygonum persicaria. Ecology 77:1791–1807.

76. Sultan, S. E. 2015. Organism and Environment: Ecological Development, Niche Construction, and Adaptation. Oxford University Press, New York, NY, United States of America.

77. Sultan, S. E. and H. G. Spencer. 2002. Metapopulation structure favors plasticity over local adaptation. Am. Nat. 160:271–283.

78. Sultan, S. E., K. Barton, and A. M. Wilczek. 2009. Contrasting patterns of transgenerational plasticity in ecologically distinct congeners. Ecology 90:1831–1839.

79. Suter L, Widmer A. 2013a. Environmental heat and salt stress induce transgenerational phenotypic changes in Arabidopsis thaliana. PLoS ONE 8:e60364. doi:10.1371/journal.pone.0060364.

80. Suter L, Widmer A. 2013b. Phenotypic effects of salt and heat stress over three generations in Arabidopsis thaliana. PLoS ONE 8:e80819. doi:10.1371/journal.pone.0080819.

81. Szendro, P. G. Vincze, A. Szasz. Pink-noise behaviour of biosystems. 2001. European Biophys. J. 30:227–231.

82. Tufto, J. 2015. Genetic evolution, plasticity, and bet-hedging as adaptive responses to temporally autocorrelated fluctuating selection: A quantitative genetic model. Evolution 69:2034–2049.

83. Uller, T. 2008. Developmental plasticity and the evolution of parental effects. Trends Ecol. Evol. 23:432–438.

84. Uller, T., S. English, and I. Pen. 2015. When is incomplete epigenetic resetting in germ cells favoured by natural selection? Proc. R. Soc. B. 282:20150682. doi: 10.1098/rspb.2015.0682.

85. Vastenhouw, N. L., K. Brunschwig, K. L. Okihara, F. Muller, M. Tijsterman, and R. H. Plasterk. 2006. Gene expression: long-term gene silencing by RNAi. Nature 442:882.

86. Verhoeven, K. J. F. and T. P. van Gurp. 2012. Transgenerational effects of stress exposure on offspring phenotypes in apomictic dandelion. PLoS ONE 7:e38605.

87. Walsh, M. R., T. Castoe, J. Holmes, M. Packer, K. Biles, M. Walsh, S. B. Munch, and D. M. Post. 2016. Local adaptation in transgenerational responses to predators. Proc. R. Soc. B. 283. 20152271. doi: 10.1098/rspb.2015.2271.

88. Whittle, C., S. Otto, M. Johnston, and J. Krochko. 2009. Adaptive epigenetic memory of ancestral temperature regime in Arabidopsis thaliana. Botany 87:650–657.

89. Wibowo, A., C. Becker, G. Marconi, J. Durr, J. Price, J. Hagmann, R. Papareddy, H. Putra, J. Kageyama, J. Becker, D. Weigel, and J. Gutierrez-Marcos. 2016. Hyperosmotic stress memory in Arabidopsis is mediated by distinct epigenetically labile sites in the genome and is restricted in the male germline by DNA glycosylase activity. Elife 5:e13546. doi: 10.7554/eLife.13546.

90. Yang, S., Z. Li, J. Yu, X. Hu, W. Dong, & S. He. 2018. El Ñino Southern Oscillation and its Impact in the Changing Climate. National Science Review doi:10.1090/nsr/nwy046.

91. Yin, J., M. Zhou, Z. Lin, Q. Li, & Y. Zhang. 2019. Transgenerational effects benefit offspring across diverse environments: a meta-analysis in plants and animals. Ecology Letters doi:10.1111/ele.13373.

92. Banta, J., D. Cruzan, M. Pigliucci. 2007. Evidence of local adaptation to coarse-grained environmental variation in *Arabidopsis thaliana*. Evolution 61:2419–2432. doi: 10.1111/j.1558-5646.2007.00189.x.

